# *InterARTIC:* an interactive web application for whole-genome nanopore sequencing analysis of SARS-CoV-2 and other viruses

**DOI:** 10.1101/2021.04.21.440861

**Authors:** James M. Ferguson, Hasindu Gamaarachchi, Thanh Nguyen, Alyne Gollon, Stephanie Tong, Chiara Aquilina-Reid, Rachel Bowen-James, Ira W. Deveson

## Abstract

**Motivation:** *InterARTIC* is an interactive web application for the analysis of viral whole-genome sequencing (WGS) data generated on Oxford Nanopore Technologies (ONT) devices. A graphical interface enables users with no bioinformatics expertise to analyse WGS experiments and reconstruct consensus genome sequences from individual isolates of viruses, such as SARS-CoV-2. *InterARTIC* is intended to facilitate widespread adoption and standardisation of ONT sequencing for viral surveillance and molecular epidemiology.

**Worked example:** We demonstrate the use of *InterARTIC* for the analysis of ONT viral WGS data from SARS-CoV-2 and Ebola virus, using a laptop computer or the internal computer on an ONT GridION sequencing device. We showcase the intuitive graphical interface, workflow customisation capabilities and job-scheduling system that facilitate execution of small- and large-scale WGS projects on any common virus.

**Implementation:** *InterARTIC* is a free, open-source web application implemented in Python. The application can be downloaded as a set of pre-compiled binaries that are compatible with all common Ubuntu distributions, or built from source. For further details please visit: https://github.com/Psy-Fer/interARTIC/.

## INTRODUCTION

Viral whole-genome sequencing (WGS) has become a critical tool used to guide local, national and international public health responses to the ongoing COVID-19 pandemic, as well as Ebola, Zika, Dengue and other viral epidemics (Eden et al. 2020; Fauver et al. 2020; Gonzalez-Reiche et al. 2020; Gudbjartsson et al. 2020; Lu et al. 2020; Meredith et al. 2020; Msomi et al. 2020; du Plessis et al. 2021; Quick et al. 2016, 2017; Rockett et al. 2020; Stubbs et al. 2020). Viral WGS is used to define the phylogenetic structure of disease outbreaks in order to better understand geographical spread, resolve transmission networks and infer the origin of unknown cases (Eden et al. 2020; Fauver et al. 2020; Gonzalez-Reiche et al. 2020; Rockett et al. 2020). Viral WGS can also identify novel strains, and monitor virus evolution over time or in response to public health interventions, such as vaccines (du Plessis et al. 2021; Fiorentini et al. 2021; Tegally et al. 2021; Williams and Burgers 2021).

Oxford Nanopore Technologies (ONT) is emerging as a leading sequencing platform for viral WGS. ONT devices are cheap, require minimal supporting laboratory infrastructure or technical expertise for sample preparation, and can be used to perform rapid viral WGS with high consensus accuracy (Bull et al. 2020). ONT’s pocket-sized MinION device can be used to perform portable viral sequencing in the field or the clinic (Quick et al. 2016, 2017; Samarakoon et al. 2020). Meanwhile, ONT’s benchtop GridION and PromethION devices enable high-throughput sequencing of large sample cohorts, with internal computers that can be utilised for data processing.

The ARTIC network for viral surveillance and molecular epidemiology has been instrumental in driving the adoption of ONT sequencing for viral WGS. ARTIC is an international consortium that has developed standardised protocols for WGS analysis of multiple common viruses, [including SARS-CoV-2](https://artic.network/ncov-2019). ARTIC researchers have also developed best-practice, open-source [bioinformatics workflows](https://github.com/artic-network/artic-ncov2019) for the reconstruction of consensus viral genome sequences from ONT sequencing data. While these command-line driven workflows are a critical resource for the community, they require specialist bioinformatics expertise that is not available to many research and public health laboratories, particularly in under-resourced areas. Moreover, the dependency of ARTIC pipelines on multiple third-party software packages creates difficulties during installation and can prevent version standardisation between different labs/sites.

To address these issues, we have developed *InterARTIC*, an interactive local web application for the analysis of viral WGS data generated on ONT devices. *InterARTIC* incorporates best-practice bioinformatics workflows for viral WGS analysis into an intuitive graphical interface that allows novice users to reconstruct consensus genome sequences from individual viral isolates, including SARS-CoV-2. The workflows are fully customisable, ensuring compatibility with different viruses and/or upstream laboratory preparations and an intelligent job-scheduling system facilitates efficient handling of large projects.

## RESULTS

### Software implementation

*InterARTIC* is a local (offline) web application for viral genomics analysis. The central component is a simple and intuitive graphical user interface developed using Python, HTML, CSS and JavaScript that encapsulates current best-practice command-line bioinformatics pipelines from ARTIC (**Fig**.**1A**). *InterARTIC* uses Flask as the web framework and features an asynchronous task queue implemented using Celery with Redis as the database backend. *InterARTIC* is provided as a standalone package that bundles the aforementioned interface, the ARTIC pipeline ([fieldbioinformatics toolkit](https://github.com/artic-network/fieldbioinformatics)) and all the dependencies.

*InterARTIC* requires no installation of third-party software and itself can be downloaded as a set of pre-compiled binaries that has been tested with Ubuntu 14, 16, 18, 20 (native Ubuntu and Ubuntu installed on Windows via windows subsystem for linux) and CentOS 7. Advanced users can alternatively build *InterARTIC* from source.

The application can execute all analysis on a standard laptop or desktop PC, after receiving sequencing data from an ONT device, such as a MinION. Alternatively, *InterARTIC* can run the analysis on the internal computer of a GridION/PromethION benchtop sequencer, generating all outputs without the need for data transfer. Importantly, *InterARTIC* was implemented with a novel software compilation/containment strategy that prevents potential conflicts with software on a user’s PC or ONT device (**Supplementary Note 1**).

### Launching the application

To use *InterARTIC*, the binaries should be downloaded, extracted and launched by running the following simple commands in a bash terminal:

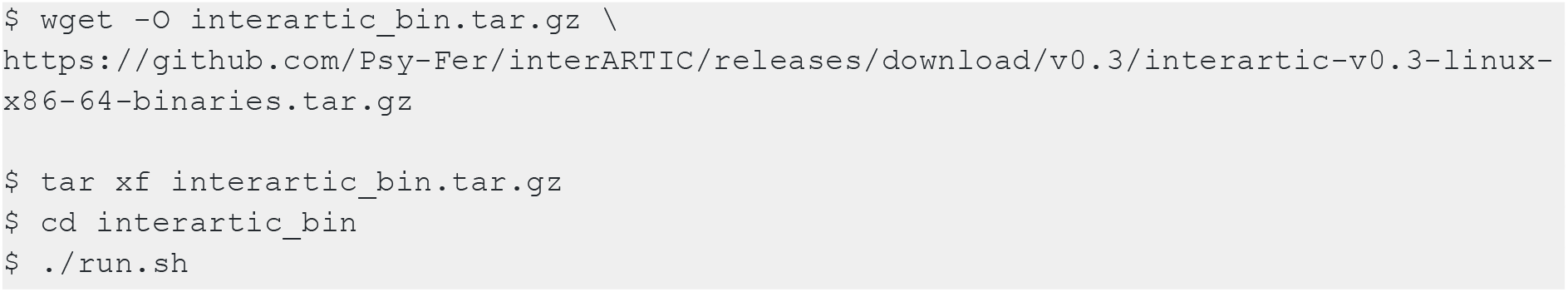

This launches an interactive session that can be accessed through a standard web browser, via a link that is printed to the standard output (by default: http://127.0.0.1:5000), as follows:

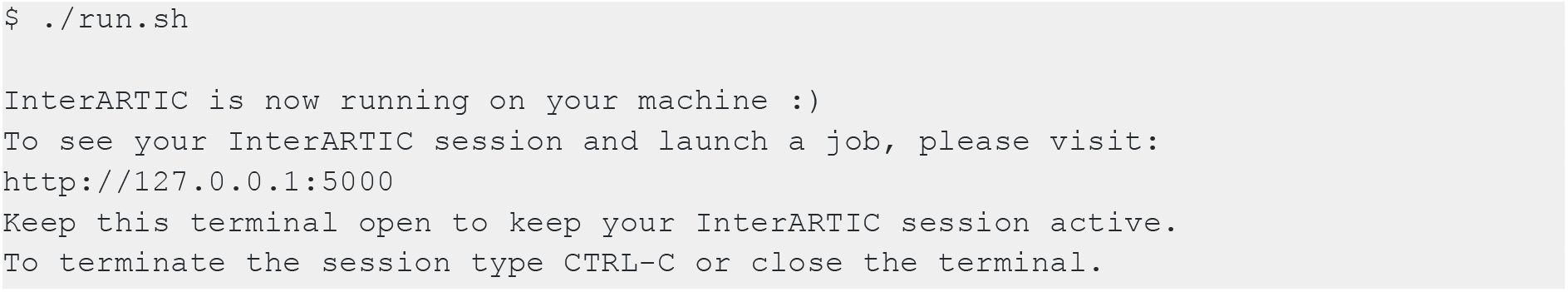

After navigating to the URL, the user can interact with the graphical interface in order to set up and launch their analysis (**Fig**.**1B**,**C**). Note that, while the user interacts with *InterARTIC* through their web browser (via localhost), all analysis is performed securely on the local hardware, where the application is hosted, and no external internet connection is required.

This article is accompanied by a simple video tutorial that helps guide new users in how to set up and operate *InterARTIC* (see **Supplementary Video 1**).

### Workflow summary

*InterARTIC* analyses amplicon-based viral WGS data generated on ONT devices. It is compatible with any virus genome, and any standard or custom amplicon primer scheme. *InterARTIC* inputs base-called sequencing reads (FASTQ format), which are typically generated during a sequencing run, using the *Guppy* base-caller within ONT’s *MinKNOW* sequencing manager software (**Fig. 1Ai**). If multiple samples were sequenced on a single flow cell, *InterARTIC* uses *Porechop* to demultiplex the library into individual sample barcodes, then runs each sample separately through the analytic workflow (**Fig. 1Aii**).

**Figure 1.**
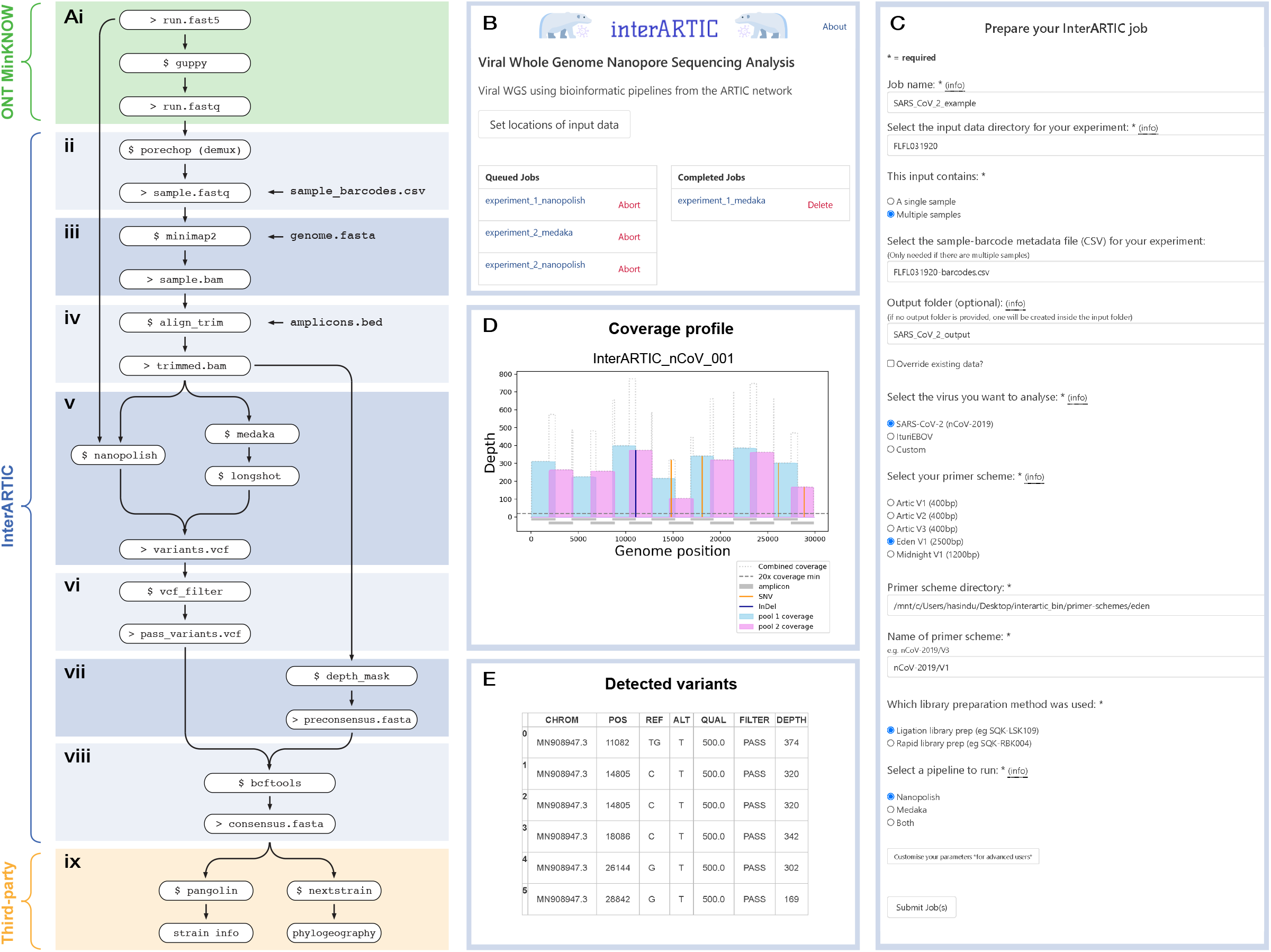
Overview of the *InterARTIC* workflow and graphical interface. (**A**) Schematic summary of a typical workflow for the analysis of ONT viral genome sequencing data. Workflow steps ii-viii (blue tiles) are executed by *InterARTIC*. (**B**) Example screenshot of the *InterARTIC* interface hompage, with queued and completed jobs displayed. (**C**) Example of the project setup page within the interface, where the user may customise their workflow. Note: full parameter selection options for advanced users are hidden. (**D**) Example of a coverage profile plot for SARS-CoV-2 that is generated and displayed by *InterARTIC*. (**E**) Example of the variant detection table that is generated and displayed.

*InterARTIC* first uses *Minimap2* to align sequencing reads to a viral reference genome (e.g., MN908947 for SARS-CoV-2; **Fig. 1Aiii**). Primer sequences are then trimmed, and alignments are down-sampled to a maximum coverage-depth threshold that can be specified by the user (**Fig. 1Aiv**). The genome (FASTA format) and primer site (BED format) reference files used during these steps can be selected from a range of popular options or supplied as custom files, according to the user’s needs. Genetic variants (SNVs and indels) are then identified relative to the reference genome. *InterARTIC* currently supports two alternative workflow choices for variant detection that utilise the popular tools *Nanopolish* or *Medaka*/*Longshot*, with the user selecting either or both workflows (**Fig. 1Av**). Low-quality variant candidates are then excluded (**Fig. 1Avi**) and regions of low coverage are masked from the reference genome (**Fig. 1Avii**). Finally, variant candidates are incorporated into the masked reference genome using *Bcftools* to generate a consensus genome sequence for the viral isolate under analysis (**Fig. 1Aviii**). This consensus genome sequence is suitable for downstream applications including lineage classification and phylogeographic analysis (**Fig. 1Aix**), and ready for upload to public repositories (e.g., [GISAID](https://www.gisaid.org/)).

In addition to the detected variants (VCF format) and consensus genome (FASTA format) output files from the workflow, *InterARTIC* also generates and displays several informative summaries and data plots that facilitate quick inspection of the data (**Fig. 1D**,**E**).

*InterARTIC* employs an asynchronous job queue system with [Celery](https://docs.celeryproject.org/en/stable/) to allow multiple jobs of the same, or differing virus and/or amplicon scheme, to be launched to allow for efficient use of compute resources (**Fig. 1B**). This is especially important if running analyses on a GridION/PromethION benchtop device, alongside other sequencing jobs.

### Example data for testing InterARTIC

To demonstrate the use of *InterARTIC*, we analysed publicly available nanopore WGS datasets from [SARS-CoV-2](https://cloudstor.aarnet.edu.au/plus/s/srVo6NEicclqQNE/download) (10 x isolates, sequenced on an ONT GridION; see Bull et al. 2020) and [Ebola](https://cloudstor.aarnet.edu.au/plus/s/9afW5kgWy1w8ZpQ/download) (2 x isolates, sequenced on an ONT MinION; see Quick et al. 2016).

Example screenshots of the workflow configuration process for each experiment are provided in **Supplementary Figures 1 & 2, a**nd example [output files](https://cloudstor.aarnet.edu.au/plus/s/SxayJLoxhQrMqil/download) generated by *InterARTIC* are also available for download.

Both projects were analysed using *InterARTIC* on a standard laptop PC, and separately on a GridION sequencing device (**Supplementary Table 1**). Indicative run-time statistics on both devices are reported in **Supplementary Table 2**. We encourage users to download these example datasets to test *InterARTIC* on their own machine.

## DISCUSSION

With new SARS-CoV-2 strains (e.g., the ‘Indian strain’ B.1.617) emerging with increasing regularity, and showing evidence in some cases of immune escape (Garcia-Beltran et al. 2021), there is a critically important ongoing role for viral genomics in the international response to COVID19. Nanopore sequencing enables rapid, cost-effective and decentralised viral WGS and - more so than any other sequencing technology - has the potential to empower local genomic surveillance initiatives across the globe.

*InterARTIC* is intended to help facilitate this process by solving two key issues. Firstly, the application enables users with no bioinformatics experience to analyse their own nanopore sequencing data to assemble complete viral genomes from individual patient isolates. This removes a major capability barrier for research and public health teams without dedicated bioinformatic scientists on their staff. Second, *InterARTIC* requires no installation and no installation of third-party software. Installation of the multiple third party-tools (and their underlying dependencies) currently required for WGS analysis can be a major challenge, even for experienced users. During this process, dependency conflicts are frequently encountered between the various software environments on a user’s PC. *InterARTIC* employs a novel containment approach, described in **Supplementary Note 1**, that removes any risk of dependency conflicts. Therefore, the application can be easily and safely run on any PC, including the on-board computer of an ONT GridION or PromethION device, removing an important resource barrier for teams lacking access to high-performance computing infrastructure.

While *InterARTIC* was originally designed for SARS-CoV-2 genomics, it is equally suitable for the analysis of any other virus. Users can supply custom inputs specific to their own virus or amplicon scheme and, to date, *InterARTIC* has been tested with SARS-CoV-2, Ebola, Dengue, Hepatitis C and Ross River viruses. We provide *InterARTIC* as a free, open-source tool to facilitate further adoption and standardisation of ONT sequencing for viral surveillance during the ongoing COVID-19 pandemic and future disease outbreaks.

## ACKNOWLEDGMENTS

We thank Charles Foster, Ellen de Vries, Igor Stevanovski and Jillian Hammond for providing feedback on *InterARTIC*. We acknowledge the following funding support: MRFF Investigator Grant MRF1173594 (to I.W.D.) and philanthropic support from The Kinghorn Foundation.

## DECLARATIONS

J.M.F. and H.G. have previously received travel and accommodation expenses to speak at Oxford Nanopore Technologies conferences. The authors declare no other competing financial or non-financial interests.

## Supplementary Note 1. The art of snake charming

*InterARTIC* development involved the use of the Python programming language and depends on several third-party Python modules and software written predominantly in Python (e.g., *Flash, Celery, ARTIC* tools, etc). The Python ecosystem (including the language itself, in addition to Python libraries) has limited backward compatibility. As a result, Python software is often compatible only with the exact version of the Python interpreter and library versions it was developed with (sometimes specific even to the minor version level). Python virtual environments and Anaconda are designed to resolve issues related to version compatibility but - at least in our experience - software installation via these methods can be complicated, especially for novice users.

Fortunately, the Python interpreter is predominantly written in C. Generally speaking, both the C language and system libraries have good backward compatibility. For instance, GLIBC is fully backward compatible. Thus, if you compile a C program on an older Linux system (e.g., Ubuntu 14) with an older compiler (e.g., gcc 4.8) and statically link third party libraries with limited backward compatibility, while dynamically linking the basic backward compatible libraries, the compiled binary would be portable on most (if not all) modern Linux systems. Of course, x86_64 binaries will not work on ARM processors, but if you compile for an older x86_64 instruction-set, it will work on all modern x86_64 processors, thanks to the backward compatibility in processor instruction sets. ARM also benefits from a similar level of backward compatibility.

In summary, if the relevant Python interpreter, all the modules and third party software are compiled and packaged with your Python code, it will be “portable”. We call this process “snake charming”, since it prevents Python modules from biting one another. For interested developers, we provide [detailed instructions](https://psy-fer.github.io/interARTIC/snakeballing/) in the InterARTIC GitHub on how snake charming was used in the development of *InterARTIC*, and how to use this technique to improve their own tools.

## Supplementary Tables

**Supplementary Table 1.** Specifications of computers used to run example workflows.

|  | Laptop | GridION |
| --- | --- | --- |
| Model | Dell XPS | GridION Mk1 |
| Processor | Intel i7-8750H | Intel i7-7700 |
| RAM | 16 GB | 64 GB |
| Disk | 1TB NVMe SSD | 4TB NVMe SSD (RAID configuration) |
| O/S | Ubuntu 16.04.7 LTS on WSL for Windows 10 | Ubuntu 16.04.7 LTS |
| Threads used | 4 | 4 |

**Supplementary Table 2.** Indicative run-times for *InterARTIC* workflow on example viral WGS datasets.

| Set | Virus | Amplicons | Samples | Pipeline | Laptop run-time (min:sec) |  |  |  | GridION run-time (min:sec) |  |  |  |
| --- | --- | --- | --- | --- | --- | --- | --- | --- | --- | --- | --- | --- |
|  |  |  |  |  | Total | Gather | Demux | Analysis | Total | Gather | Demux | Analysis |
| 1 | SARS-CoV-2 | Eden V1 2.5kb | 10 | Nanopolish | 32:19 | 00:07 | 17:56 | 14:05 | 25:09 | 00:05 | 13:42 | 11:33 |
|  |  |  |  | Medaka-Longshot | 30:24 | 0:11 | 17:53 | 12:09 | 21:41 | 00:05 | 13:43 | 07:45 |
| 3 | Ebola | Artic V1 400bp | 2 | Nanopolish | 03:57 | 00:02 | 02:23 | 01:30 | 02:55 | 00:01 | 01:42 | 01:10 |
|  |  |  |  | Medaka-Longshot | 04:09 | 00:03 | 02:24 | 01:42 | 02:54 | 00:01 | 01:40 | 01:12 |

**Supplementary Figure 1.**
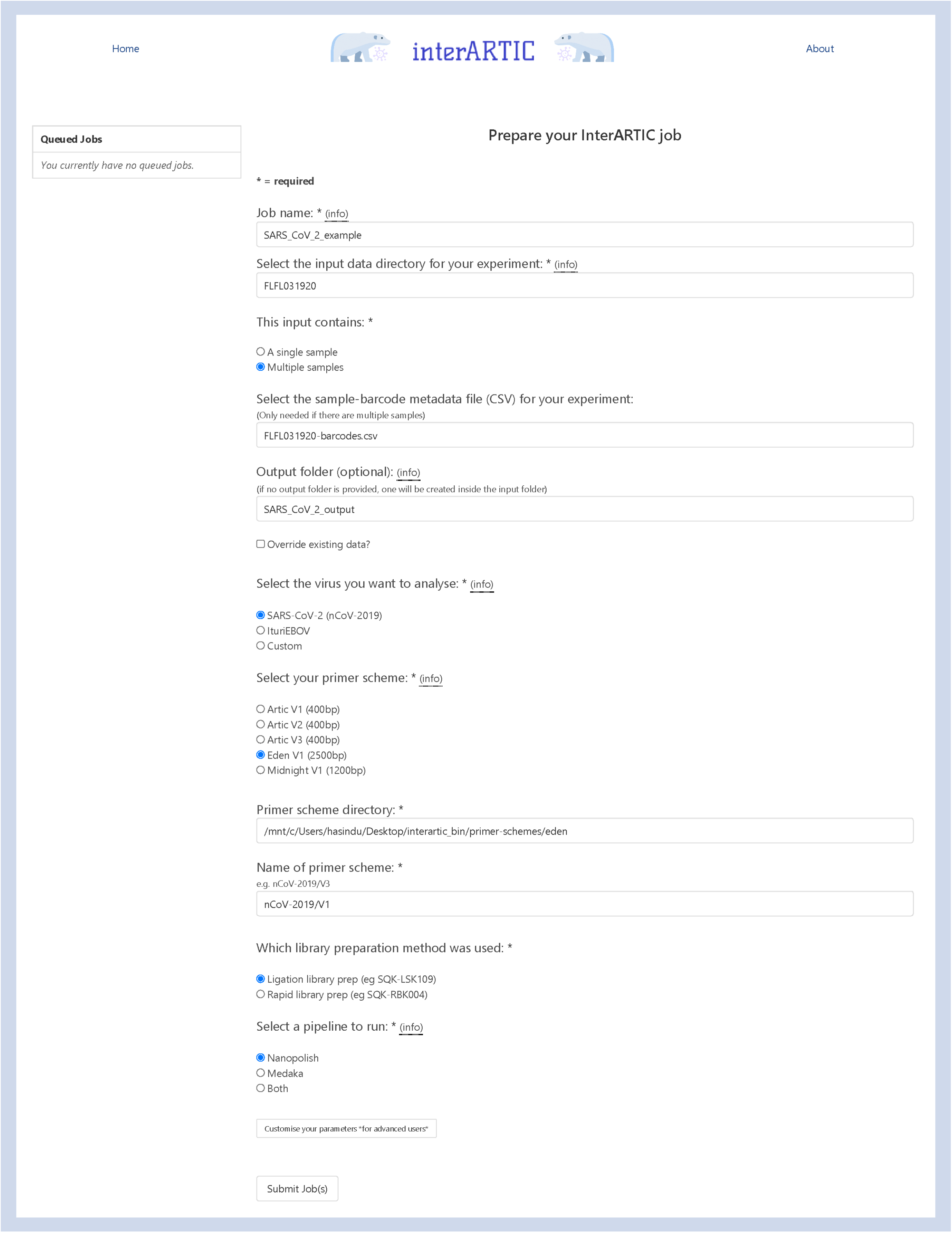
Example job configuration for *InterARTIC* analysis of SARS-CoV-2 whole-genome sequencing.

**Supplementary Figure 2.**
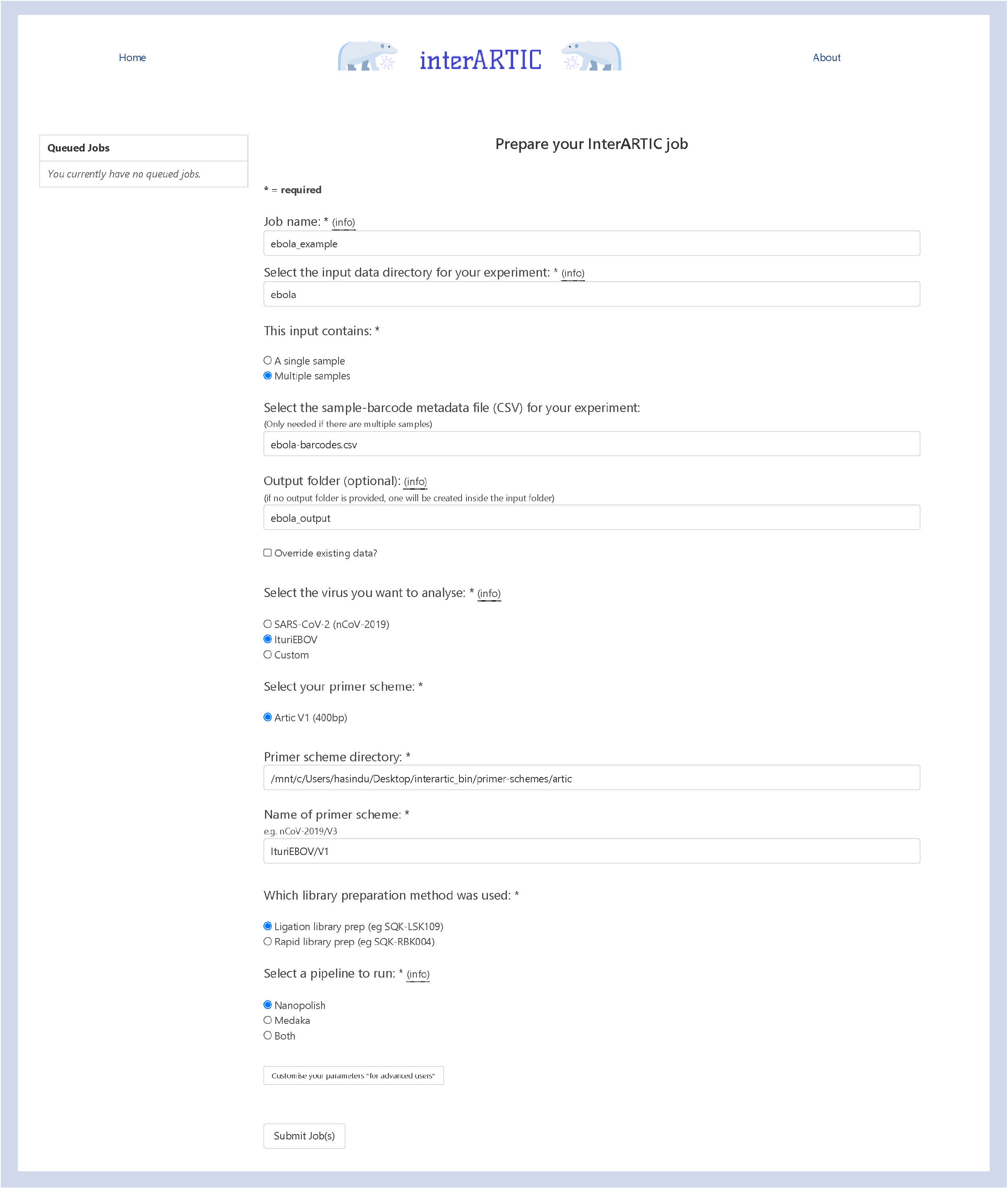
Example job configuration for *InterARTIC* analysis of Ebola whole-genome sequencing.

## REFERENCES

Bull RA, Adikari TN, Ferguson JM, Hammond JM, Stevanovski I, Beukers AG, Naing Z, Yeang M, Verich A, Gamaarachchi H, et al. 2020. Analytical validity of nanopore sequencing for rapid SARS-CoV-2 genome analysis. Nat Commun 11: 6272.

du Plessis L, McCrone JT, Zarebski AE, Hill V, Ruis C, Gutierrez B, Raghwani J, Ashworth J, Colquhoun R, Connor TR, et al. 2021. Establishment and lineage dynamics of the SARS-CoV-2 epidemic in the UK. Science 371: 708–712.

Eden J-S, Rockett R, Carter I, Rahman H, de Ligt J, Hadfield J, Storey M, Ren X, Tulloch R, Basile K, et al. 2020. An emergent clade of SARS-CoV-2 linked to returned travellers from Iran. Virus Evol 6: veaa027.

Fauver JR, Petrone ME, Hodcroft EB, Shioda K, Ehrlich HY, Watts AG, Vogels CBF, Brito AF, Alpert T, Muyombwe A, et al. 2020. Coast-to-Coast Spread of SARS-CoV-2 during the Early Epidemic in the United States. Cell 181: 990-996.e5.

Fiorentini S, Messali S, Zani A, Caccuri F, Giovanetti M, Ciccozzi M, Caruso A. 2021. First detection of SARS-CoV-2 spike protein N501 mutation in Italy in August, 2020. Lancet Infect Dis. http://dx.doi.org/10.1016/S1473-3099(21)00007-4.

Garcia-Beltran WF, Lam EC, St Denis K, Nitido AD, Garcia ZH, Hauser BM, Feldman J, Pavlovic MN, Gregory DJ, Poznansky MC, et al. 2021. Multiple SARS-CoV-2 variants escape neutralization by vaccine-induced humoral immunity. Cell 184: 2523.

Gonzalez-Reiche AS, Hernandez MM, Sullivan MJ, Ciferri B, Alshammary H, Obla A, Fabre S, Kleiner G, Polanco J, Khan Z, et al. 2020. Introductions and early spread of SARS-CoV-2 in the New York City area. Science 369: 297–301.

Gudbjartsson DF, Helgason A, Jonsson H, Magnusson OT, Melsted P, Norddahl GL, Saemundsdottir J, Sigurdsson A, Sulem P, Agustsdottir AB, et al. 2020. Spread of SARS-CoV-2 in the Icelandic Population. N Engl J Med 382: 2302–2315.

Lu J, du Plessis L, Liu Z, Hill V, Kang M, Lin H, Sun J, François S, Kraemer MUG, Faria NR, et al. 2020. Genomic Epidemiology of SARS-CoV-2 in Guangdong Province, China. Cell 181: 997-1003.e9.

Meredith LW, Hamilton WL, Warne B, Houldcroft CJ, Hosmillo M, Jahun AS, Curran MD, Parmar S, Caller LG, Caddy SL, et al. 2020. Rapid implementation of SARS-CoV-2 sequencing to investigate cases of health-care associated COVID-19: a prospective genomic surveillance study. Lancet Infect Dis 20: 1263–1271.

Msomi N, Mlisana K, de Oliveira T, Network for Genomic Surveillance in South Africa writing group. 2020. A genomics network established to respond rapidly to public health threats in South Africa. Lancet Microbe 1: e229–e230.

Quick J, Grubaugh ND, Pullan ST, Claro IM, Smith AD, Gangavarapu K, Oliveira G, Robles-Sikisaka R, Rogers TF, Beutler NA, et al. 2017. Multiplex PCR method for MinION and Illumina sequencing of Zika and other virus genomes directly from clinical samples. Nat Protoc 12: 1261–1276.

Quick J, Loman NJ, Duraffour S, Simpson JT, Severi E, Cowley L, Bore JA, Koundouno R, Dudas G, Mikhail A, et al. 2016. Real-time, portable genome sequencing for Ebola surveillance. Nature 530: 228–232.

Rockett RJ, Arnott A, Lam C, Sadsad R, Timms V, Gray K-A, Eden J-S, Chang S, Gall M, Draper J, et al. 2020. Revealing COVID-19 transmission in Australia by SARS-CoV-2 genome sequencing and agent-based modeling. Nat Med 26: 1398–1404.

Samarakoon H, Punchihewa S, Senanayake A, Hammond JM, Stevanovski I, Ferguson JM, Ragel R, Gamaarachchi H, Deveson IW. 2020. Genopo: a nanopore sequencing analysis toolkit for portable Android devices. Commun Biol 3: 538.

Stubbs SCB, Blacklaws BA, Yohan B, Yudhaputri FA, Hayati RF, Schwem B, Salvaña EM, Destura RV, Lester JS, Myint KS, et al. 2020. Assessment of a multiplex PCR and Nanopore-based method for dengue virus sequencing in Indonesia. Virol J 17: 24.

Tegally H, Wilkinson E, Giovanetti M, Iranzadeh A, Fonseca V, Giandhari J, Doolabh D, Pillay S, San EJ, Msomi N, et al. 2021. Emergence of a SARS-CoV-2 variant of concern with mutations in spike glycoprotein. Nature. http://dx.doi.org/10.1038/s41586-021-03402-9.

Williams TC, Burgers WA. 2021. SARS-CoV-2 evolution and vaccines: cause for concern? Lancet Respir Med. http://dx.doi.org/10.1016/S2213-2600(21)00075-8.

